# The structural basis for kinetochore stabilization by Cnn1/CENP-T

**DOI:** 10.1101/2020.05.04.077578

**Authors:** Stephen M. Hinshaw, Stephen C. Harrison

**Affiliations:** Harvard Medical School, Howard Hughes Medical Institute

## Abstract

Chromosome segregation depends on a regulated connection between spindle microtubules and centromeric DNA. The kinetochore, a massive modular protein assembly, mediates this connection and also serves as a signaling hub that integrates and responds to changing cues during the cell cycle. Kinetochore functions evolve as the cell cycle progresses, culminating in the assurance of a persistent chromosome-microtubule connection during anaphase, when sister chromatids must transit into daughter cells uninterrupted. We previously determined the structure of the Ctf19 complex, a group of kinetochore proteins at the centromeric base of the kinetochore. We now present a high-resolution structure of a Ctf19 complex sub-assembly involved in centromere-microtubule contact: the Ctf3 complex bound to the Cnn1-Wip1 heterodimer. The resulting composite model of the Ctf19 complex and live-cell imaging experiments provide a mechanism for Cnn1-Wip1 recruitment to the kinetochore. The mechanism suggests feedback regulation of Ctf19 complex assembly and unanticipated similarities in kinetochore organization between yeast and vertebrates.

## INTRODUCTION

The kinetochore controls chromosome segregation by coordinating chromosome-microtubule contact with the events of the cell division cycle. There are two classes of kinetochore proteins: those that assemble on centromeric DNA (the inner kinetochore) and those that make up the microtubule interaction apparatus (the outer kinetochore). The enzymes that drive the cell cycle affect the two functional classes accordingly. For example, chromosome misalignment generates a signal that causes the DNA-bound kinetochore assembly platform to solidify while weakening the connection between the microtubule and its interacting proteins (Akiyoshi et al., 2013; Cheeseman et al., 2006; DeLuca et al., 2006; Kim and Yu, 2015; Pinsky et al., 2006). During anaphase, when microtubules pull sister chromatids to opposite spindle poles, outer kinetochore proteins are enriched, ensuring a persistent connection to the depolymerizing filament (Dhatchinamoorthy et al., 2017; Gascoigne and Cheeseman, 2013).

Most inner kinetochore proteins belong to a large group of factors that associate biochemically. The group is called the Ctf19 complex (Ctf19c) in budding yeast and the Constitutive Centromere Associated Network (CCAN) in vertebrates (Cheeseman et al., 2002; De Wulf et al., 2003; Foltz et al., 2006; Okada et al., 2006). Although the structures and identities of these proteins are mostly conserved, their presence varies across eukaryotic species (Plowman et al., 2019). Nevertheless, all 13 yeast factors assemble into an interdigitated structure thought to embrace a centromere-defining nucleosome containing the histone H3 variant Cse4 (CENP-A in vertebrates) (Hinshaw and Harrison, 2019). When isolated by means of an affinity-tagged basal factor (Ame1/CENP-U), Ctf19c proteins have a strictly hierarchical mode of association such that outer factors require inner factors for their recruitment to the complex (Pekgoz Altunkaya et al., 2016). This observation led to a model for Ctf19c assembly mostly consistent with published cell biology experiments and with the recent structural information.

Among Ctf19c sub-complexes, the outermost is a five-protein module containing the Ctf3 heterotrimer (Ctf3c, CENP-H/I/K in vertebrates) and the Cnn1-Wip1 heterodimer (CENP-T/W in vertebrates, **Figure 1A, Table 1**). When the Ctf19c is purified from extract, Ctf3c proteins are required for Cnn1-Wip1 recovery, while the reverse is not true (Pekgoz Altunkaya et al., 2016). Ctf3 itself is required in cells for recruitment of the replicative kinase DDK and the SUMO protease Ulp2 (Hinshaw et al., 2017; Suhandynata et al., 2019). The molecular details of DDK and Ulp2 recruitment are not yet clear. The mechanism of Ctf3c recruitment to the kinetochore is also still unclear, as yeast strains lacking Ctf19 show reduced but not ablated Ctf3-GFP kinetochore localization (Pot et al., 2003). Thus, the biochemical experiments performed with cell extracts and described above do not completely recapitulate the logic of kinetochore proteins were purified centromeric Ctf19c assembly.

**Table 1.** Ctf3c and Cnn1-Wip1 proteins in different species

| *S. cerevisiae* | *S. pombe* | *H. sapiens* |
| --- | --- | --- |
| Ctf3 | Mis6 | CENP-I |
| Mcm16 | Fta3 | CENP-H |
| Mcm22 | Sim4 | CENP-K |
|  |  | CENP-M |
| Cnn1 | Cnp20 | CENP-T |
| Wip1 | Wip1 | CENP-W |
| Mhf1* | Mhf1 | CENP-S |
| Mhf2* | Mhf2 | CENP-X |

\* No association with Ctf19c factors *in vitro* (Pekgoz Altunkaya et al., 2016)

**Table 2.**
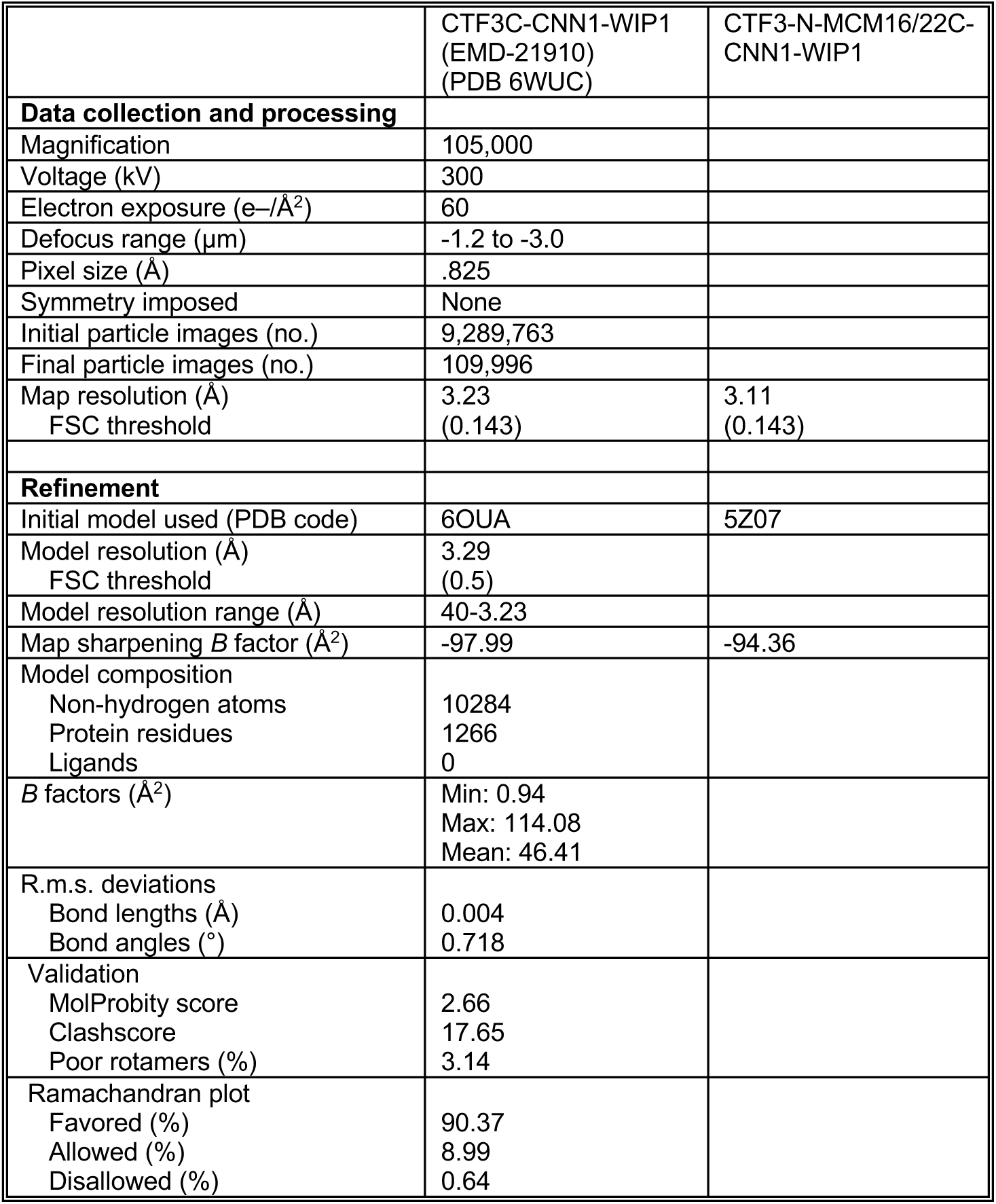
Cryo-EM data collection, refinement, and validation

|  | CTF3C-CNN1-WIP1<br>(EMD-21910)<br>(PDB 6WUC) | CTF3-N-MCM16/22C-CNN1-WIP1 |
| --- | --- | --- |
| **Data collection and processing** |  |  |
| Magnification | 105,000 |  |
| Voltage (kV) | 300 |  |
| Electron exposure (e-/Å<sup>2</sup>) | 60 |  |
| Defocus range (μm) | -1.2 to -3.0 |  |
| Pixel size (Å) | .825 |  |
| Symmetry imposed | None |  |
| Initial particle images (no.) | 9,289,763 |  |
| Final particle images (no.) | 109,996 |  |
| Map resolution (Å) | 3.23 | 3.11 |
| FSC threshold | (0.143) | (0.143) |
| **Refinement** |  |  |
| Initial model used (PDB code) | 6OUA | 5Z07 |
| Model resolution (Å) | 3.29 |  |
| FSC threshold | (0.5) |  |
| Model resolution range (Å) | 40-3.23 |  |
| Map sharpening *B* factor (Å<sup>2</sup>) | -97.99 | -94.36 |
| Model composition |  |  |
| Non-hydrogen atoms | 10284 |  |
| Protein residues | 1266 |  |
| Ligands | 0 |  |
| *B* factors (Å<sup>2</sup>) | Min: 0.94<br>Max: 114.08<br>Mean: 46.41 |  |
| R.m.s. deviations |  |  |
| Bond lengths (Å) | 0.004 |  |
| Bond angles (°) | 0.718 |  |
| Validation |  |  |
| MolProbity score | 2.66 |  |
| Clashscore | 17.65 |  |
| Poor rotamers (%) | 3.14 |  |
| Ramachandran plot |  |  |
| Favored (%) | 90.37 |  |
| Allowed (%) | 8.99 |  |
| Disallowed (%) | 0.64 |  |

**Table 3.** Model information

| PROTEIN | CHAIN ID | CHAIN LENGTH (TOTAL; MODELED) | TEMPLATE | PROCEDURE | MODELLED RESIDUES |
| --- | --- | --- | --- | --- | --- |
| Mcm16 | H | 181; 179 | 6OUA, 5Z07 | Modify, build *de novo* | 3-181 |
| Ctf3 | I | 733; 712 | 6OUA, 5Z07 | Modify, build *de novo* | 0-268, 276-317, 327-728 |
| Mcm22 | K | 239; 235 | 6OUA, 5Z07 | Modify, build *de novo* | 5-239 |
| Cnn1 | T | 361; 87 | 3B0C | Modify | 274-317, 322-361 |
| Wip1 | W | 89; 57 | 3B0C | Modify | 15-25, 36-68, 77-87 |

**Table 4.** Yeast strains used in this study

| STRAIN NUMBER | GENOTYPE | REFERENCE |
| --- | --- | --- |
| SMH75 | *MATa CTF3-GFP::HisMX SPC110-mCherry::hphMX* | (Huh et al., 2003) |
| SMH728 | *MATa CTF3-GFP::HisMX SPC110-mCherry::hphMX MCM22-3FLAG::KanMX* | This study |
| SMH746 | *MATa CTF3-GFP::HisMX SPC110-mCherry::hphMX MCM22-3FLAG::KanMX iml3Δ::natMX* | This study |
| SMH767 | *MATa CTF3-GFP::HisMX SPC110-mCherry::hphMX ctf19Δ::KanMX* | This study |
| SMH768 | *MATa CTF3-GFP::HisMX SPC110-mCherry::hphMX cnn1Δ::KanMX* | This study |
| SMH749 | *MATa CTF3-GFP::HisMX SPC110-mCherry::hphMX mcm22-2A::KanMX* | This study |
| SMH783 | *MATa CTF3-GFP::HisMX SPC110-mCherry::hphMX ctf19Δ::KanMX cnn1Δ::KanMX* | This study |
| SMH784 | *MATa CTF3-GFP::HisMX SPC110-mCherry::hphMX ctf19Δ::KanMX mcm22-2A::KanMX* | This study |

W303 derivatives:
| STRAIN NUMBER | GENOTYPE | REFERENCE |
| --- | --- | --- |
| PDW2002 | *MATa CNN1-3GFP::KanMX SPC110-mCherry::hphMX* | (Bock et al., 2012) |
| SMH798 | *MATa CNN1-3GFP::KanMX SPC110-mCherry::hphMX MCM22-3FLAG::HisMX* | This study |
| SMH805 | *MATa CNN1-3GFP::KanMX SPC110-mCherry::hphMX mcm22Δ::HisMX* | This study |
| SMH809 | *MATa CNN1-3GFP::KanMX SPC110-mCherry::hphMX mcm22Δ::HisMX* | This study |
| SMH799 | *MATa CNN1-3GFP::KanMX SPC110-mCherry::hphMX mcm22-2A::HisMX* | This study |
| SMH800 | *MATa CNN1-3GFP::KanMX SPC110-mCherry::hphMX mcm22-2A::HisMX* | This study |

**Table 5.** Plasmids used in this study

| PLASMID NUMBER | GENOTYPE | REFERENCE |
| --- | --- | --- |
| pSMH145 | pLIC-Tra His6-TEV-Ctf3; His6-TEV-Mcm16; His6-TEV-Mcm22 | (Hinshaw et al., 2017) |
| pSMH1712 | pLIC-Tra His6-TEV-Ctf3; His6-TEV-Mcm16; His6-TEV-Mcm22-206,210A (Ctf3c-2A mutant) | This study |
| pSMH1193 | pLICBac Cnn1 (untagged) | (Hinshaw and Harrison, 2019) |
| pSMH1198 | pLICBac His6-Wip1 | (Hinshaw and Harrison, 2019) |
| pSMH1693 | pFA6a-MCM22-6Gly-3FLAG-KanMX6 (BsiWI/Sall) | This study |
| pSMH1709 | pFA6a-mcm22-2A-6Gly-3FLAG-KanMX6 (BsiWI/Sall) | This study |
| pSMH1776 | pFA6a-MCM22-6Gly-3FLAG-HisMX (BsiWI/Sall) | This study |
| pSMH1777 | pFA6a-mcm22-2A-6Gly-3FLAG- HisMX (BsiWI/Sall) | This study |

**Figure 1.**
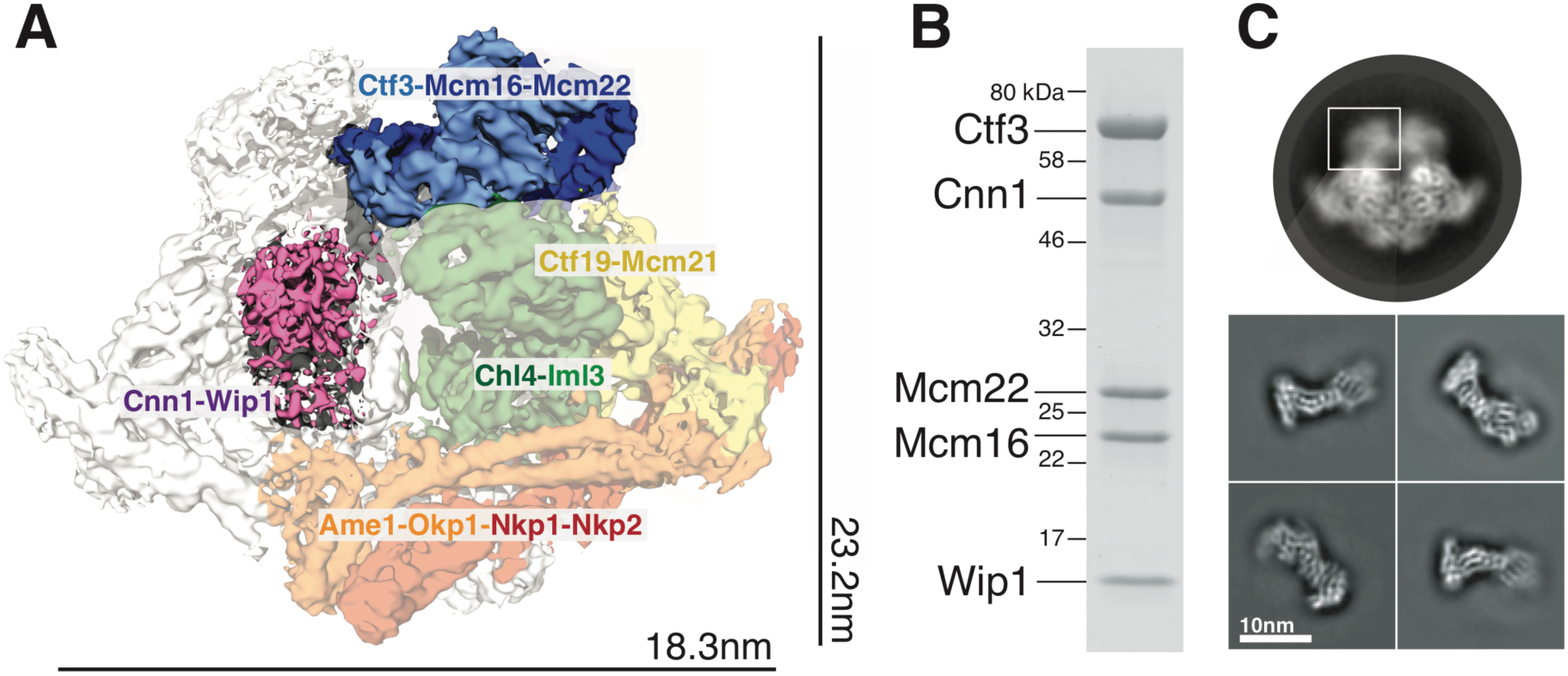
Reconstitution of the Ctf3c bound to Cnn1-Wip1. **(A)** The structure of the dimeric Ctf19c with the five-protein Ctf3c-Cnn1-Wip1 complex highlighted. The Ctf19c density shows two copies of each protein subunit, and the unique protomer is colored accord to subcomplex identity. **(B)** Ctf3c-Cnn1-Wip1 sample used for structure determination (SDS-PAGE). **(C)** A representative two-dimensional class average showing the intact Ctf19c with the region corresponding to Ctf3c-Cnn1-Wip1 highlighted (top). Images below show two-dimensional class averages from the purified Ctf3c-Cnn1-Wip1 sample shown in (B).

Cnn1 and Wip1 heterodimerize through complementary C-terminal histone folds. Their vertebrate homologs bind and bend DNA (along with partners CENP-S/X) (Hori et al., 2008; Takeuchi et al., 2014), and key CENP-T DNA-binding residues are conserved in yeast Cnn1 (Takeuchi et al., 2014). Cnn1 has an extended N-terminal region (Cnn1-N, ∼270 amino acid residues) that contacts the microtubule lattice indirectly by binding the Spc24 and Spc25 components of the Ndc80 complex (Ndc80c) (Malvezzi et al., 2013; Nishino et al., 2013; Schleiffer et al., 2012). Cdc28/CDK1, Mps1, and Ipl1 all phosphorylate Cnn1-N and modulate its kinetochore recruitment (Bock et al., 2012; Thapa et al., 2015). In particular, Mps1 inhibits Ndc80c binding by phosphorylating Cnn1-S74 (Malvezzi et al., 2013). The Cnn1 histone fold contacts the Ctf3c through a histone fold extension motif (HFE) conserved among CENP-T proteins (Pekgoz Altunkaya et al., 2016). Cnn1-HFE mutants are defective in kinetochore localization, and the Cnn1-HFE and Cnn1-N both contribute to Cnn1 kinetochore localization (Thapa et al., 2015).

To better understand the cell cycle-dependent recruitment and activities of Ctf19c proteins, we determined the structure of the Ctf3c bound to the Cnn1-Wip1 heterodimer. We used the structure to identify Ctf3c mutations that disable Cnn1-Wip1 binding and studied the effects of these mutations *in vivo*. These experiments provide evidence that Cnn1 integrates mitotic kinase activity to regulate inner kinetochore assembly.

## RESULTS

### Reconstitution and cryo-EM structure of the Ctf3c-Cnn1-Wip1 complex

Recombinant Ctf3c interacts stably with Cnn1-Wip1, making a five-protein complex with one copy of each protein component (**Figure 1B**). We purified the reconstituted complex by size exclusion chromatography and determined its structure by cryo-EM. Two-dimensional class averages show projected density that matches a previous reconstruction of the Ctf3c with additional density corresponding to the N-terminal part of Ctf3, Wip1, and Cnn1 (**Figure 1C).** Images from these class averages contributed to a map resolved to a final resolution of 3.2 Å for the full complex and to 3.1 Å for Ctf3-N, Wip1, and Cnn1 (**Figure S1**).

The two HEAT repeat domains of Ctf3 define a modular Ctf3c structure (**Figure 2A**). The N-terminal HEAT repeats (Ctf3-N) contact the C-terminal part of Mcm 16/22 (Mcm16/22-C), while the C-terminal HEAT repeats (Ctf3-C) contact the N-terminal part of Mcm16/22 (Mcm16/22-N). The resolution is consistent throughout the map with the exception of the extremities along the long axis of the complex (**Figure S2**), due to subtle flexing at the midpoint observable in some three-dimensional classifications. Further refinement of the Ctf3-N module remediated this issue slightly.

**Figure 2.**
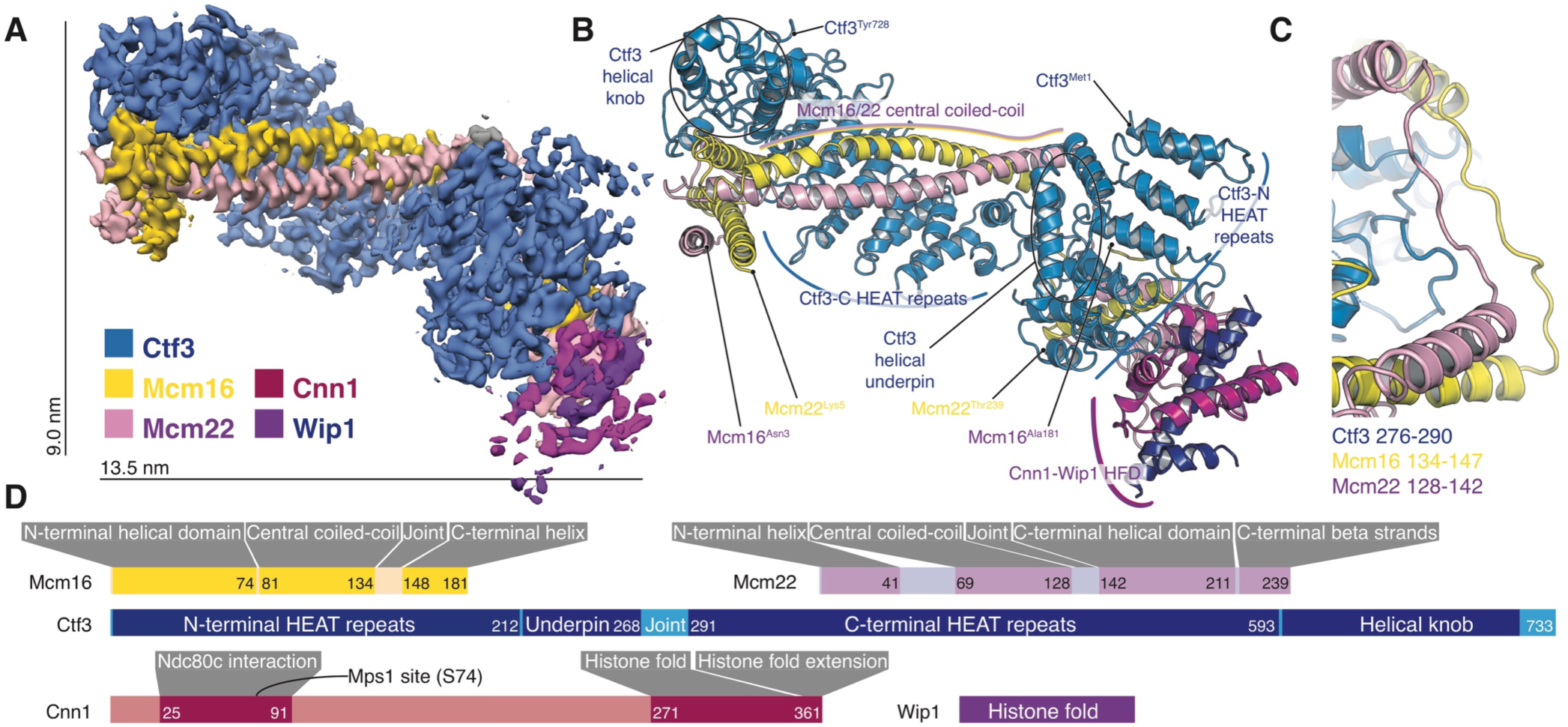
Structure of the Ctf3c bound to Cnn1-Wip1. **(A)** Cryo-EM density map showing the Ctf3c bound to Cnn1-Wip1. The map is colored according to underlying protein subunit identity. **(B)** Model of the Ctf3c bound to Cnn1-Wip1. **(C)** Close-up view of the Ctf3c joint region that connects the Ctf3-N and Ctf3-C modules. Numbers are given for amino acid residues not contributing to the helical segments shown. **(D)** Domain diagram describing the Ctf3c and Cnn1-Wip1. The Ndc80c-interacting region of Cnn1 contains the Mps1 phosphorylation site (Cnn1-S74) and corresponds to a previously-described Spc24/Spc25 interaction sequence (SIS) (Malvezzi et al., 2013; Thapa et al., 2015). Dark colors indicate ordered regions, while light colors mark segments without defined secondary structure elements, including the joint region shown in (C).

### Molecular model of the complete Ctf3c

The high-resolution cryo-EM map enables modeling of the complete Ctf3c (**Figure 2B**). Previous studies have reported molecular structures, determined by cryo-EM or crystallography, of Ctf3-C with Mcm16/22-N from *S. cerevisiae* and of Ctf3-N with Mcm16/22-C from related yeast species (Hinshaw et al., 2019; Hu et al., 2019; Zhang et al., 2020). In cryo-EM reconstructions of the complete Ctf19c, only the Ctf3-C module is well-resolved (Hinshaw and Harrison, 2019; Yan et al., 2019). The new Ctf3c model shows how these structural modules relate to each other and provides a high-resolution model of *S. cerevisiae* Ctf3-N with Mcm16/22-C. Comparison with a low-resolution model of the human CENP-H/I/K/M complex suggests the overall organization is conserved (Basilico et al., 2014).

The two Ctf3 HEAT repeat domains engage a discontinuous Mcm16/22 coiled-coil by distinct binding modes. The convex surface of Ctf3-N (Ctf3 1-223) provides a binding surface for Mcm16/22-C (Mcm16 148-181, Mcm22 142-239), while the concave inner surface supports a two-helix underpinning motif (Ctf3 228-268). The concave surface of Ctf3-C (Ctf3 290-733) binds an extended Mcm16/22-N coiled-coil (Mcm16 81-134, Mcm22 69-128), with terminal helices of Mcm16/22-N (Mcm16 1-41; Mcm22 1-74) protruding and stabilizing a Ctf3 C-terminal helical knob, as previously described (Hinshaw et al., 2019). Extended regions of Mcm16/22 lacking secondary structure but visible in the density (Mcm16 134-147, Mcm22 128-142) link the two Ctf3 HEAT modules (**Figure 2C**), which themselves are connected by an atypical HEAT repeat not visible in previous models (Ctf3 290-302).

### Observation and validation of the Ctf3c-Cnn1 interface

The Cnn1-HFE, which contacts Mcm22, is well-resolved in the new density, allowing unambiguous modeling of its interaction with the Ctf3c (**Figure 3A**). Previous work showed that contact between the Ctf3c and Cnn1 depends on a set of conserved residues in the Cnn1-HFE (Cnn1-E346, L350, E351) (Pekgoz Altunkaya et al., 2016). Cnn1-L350 satisfies a hydrophobic pocket made by Ctf3-F142, and Cnn1-E351 caps the pocket by making contact with the backbone amide of Ctf3-S143. Cnn1-E346 does not make obvious contact with the Ctf3c. Together, Mcm22-R206 and Mcm22-R210 coordinate the negative charge of the Cnn1-E357 side chain, and Cnn1-S354 further stabilizes this connection.

**Figure 3.**
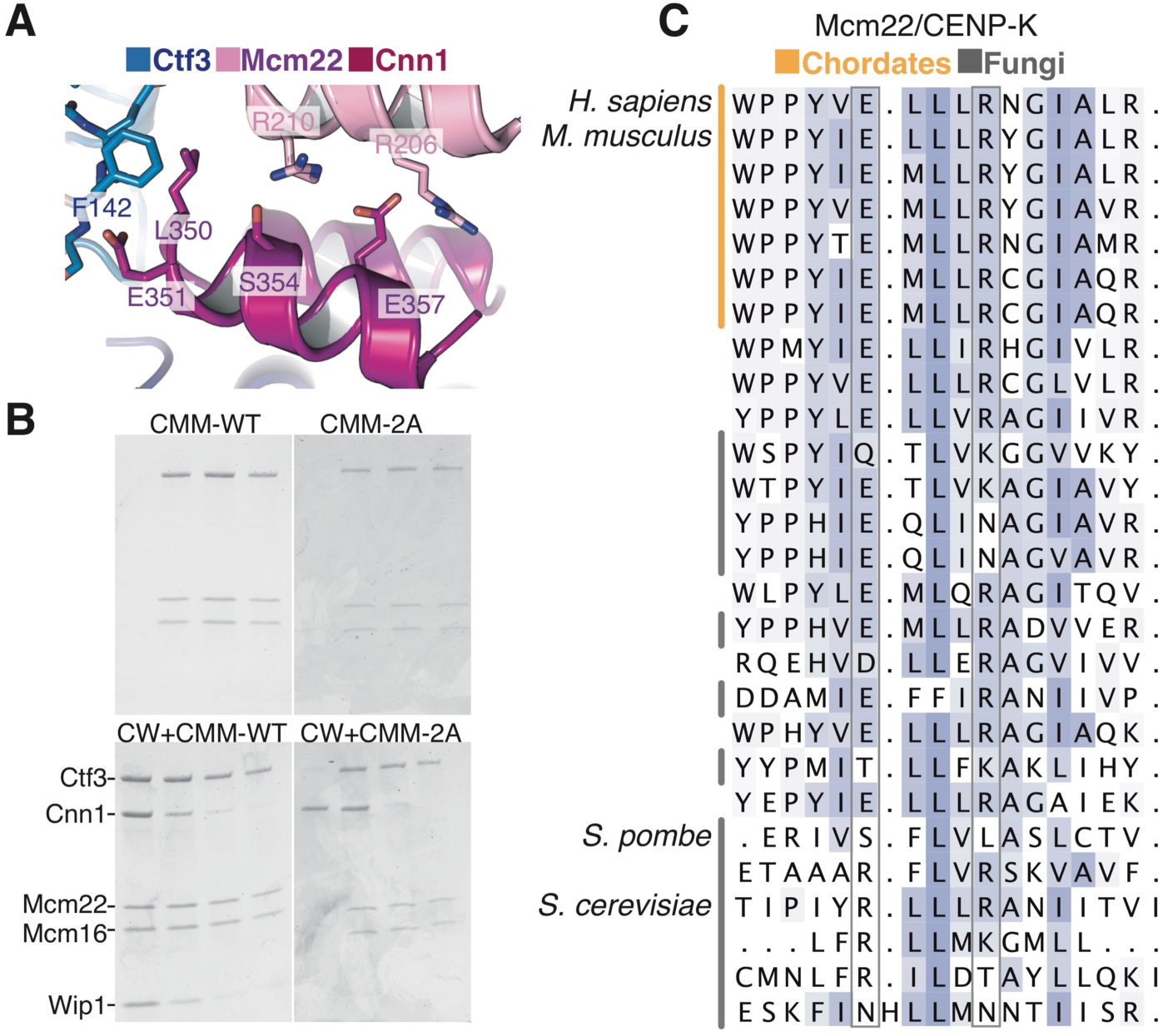
Observation and validation of the Cnn1-Ctf3c interface. **(A)** Close-up view of the Ctf3c-Cnn1 interface. The two Cnn1 helices shown constitute the Cnn1-HFE. **(B)** Ctf3c binding to Cnn1-Wip1 and perturbation by the Mcm22-2A mutations. Equivalent gel filtration fractions are shown on each gel. The top panels show recombinant Ctf3c (left) or Ctf3c containing the Mcm22-2A mutant protein (right). The results of mixing with recombinant Cnn1-Wip1 are shown in the bottom panels. **(C)** Multiple sequence alignment showing Mcm22 conservation at the Cnn1 contact site. Four species are noted, and sequences from chordates and fungi are highlighted at left. Gray boxes denote residues aligning with *S. cerevisiae* Mcm22-R206 and -R210.

The Cnn1-Wip1 histone fold domain is sufficient to interact with the Ctf3c (Pekgoz Altunkaya et al., 2016). Mutation of Mcm22-R206 and -R210 to alanine (Mcm22-2A) completely ablated this interaction (**Figure 3B**). Analysis of amino acid conservation throughout eukaryotes shows that the Cnn1 binding site is more conserved than the rest of the Ctf3c (**Figure 3C, Figure S3**). Indeed, Mcm22-R210 aligns with basic amino acids (arginine or lysine) for most species examined. Vertebrate CENP-T/W binds CENP-S/X to make a histone heterotetramer (Nishino et al., 2012). The CENP-S/X binding site would be fully exposed were CENP-T/W to engage CENP-H/I/K as it does the yeast Ctf3c. Therefore, the mode of interaction between Cnn1-Wip1 and the Ctf3c is likely maintained throughout eukaryotes.

### The Ctf3c and Cnn1-Wip1 depend on each other for kinetochore recruitment

To test the idea that Ctf19c assembly is hierarchical, we measured Cnn1 kinetochore localization in cells expressing Cnn1-3GFP (**Figure 4A**). Alignment of all measurements according to anaphase onset (determined by separation of Spc110-mCherry) shows that Cnn1 recruitment peaks in anaphase and dissipates as cells enter the cell cycle (**Figure 4B**) (Bock et al., 2012). If the Ctf3c is strictly required for Cnn1 localization, mutations that disrupt the Ctf3c-Cnn1 interaction should ablate Cnn1-GFP localization. Instead, cells expressing Cnn1-3GFP and *mcm22-2A* lack the Cnn1 kinetochore signal until anaphase, when Cnn1 intensity at kinetochores peaks but does not reach wild-type levels. These cells are indistinguishable from *mcm22Δ* cells except that, unlike *mcm22Δ* cells, they are insensitive to the Mps1 inhibitor, cincreasin (Dorer et al., 2005) (**Figure S4B**), indicating that Ctf3c functions unrelated to Cnn1-Wip1 recruitment are preserved. In a metaphase arrest, both mutants lack the robust Cnn1-3GFP kinetochore signal seen in a wild-type background (**Figure S4C**), showing that Cnn1 localization does not occur before anaphase in the mutants. Thus, the Ctf3c is required for Cnn1 localization until anaphase, when a second pathway for Cnn1 localization also contributes.

**Figure 4.**
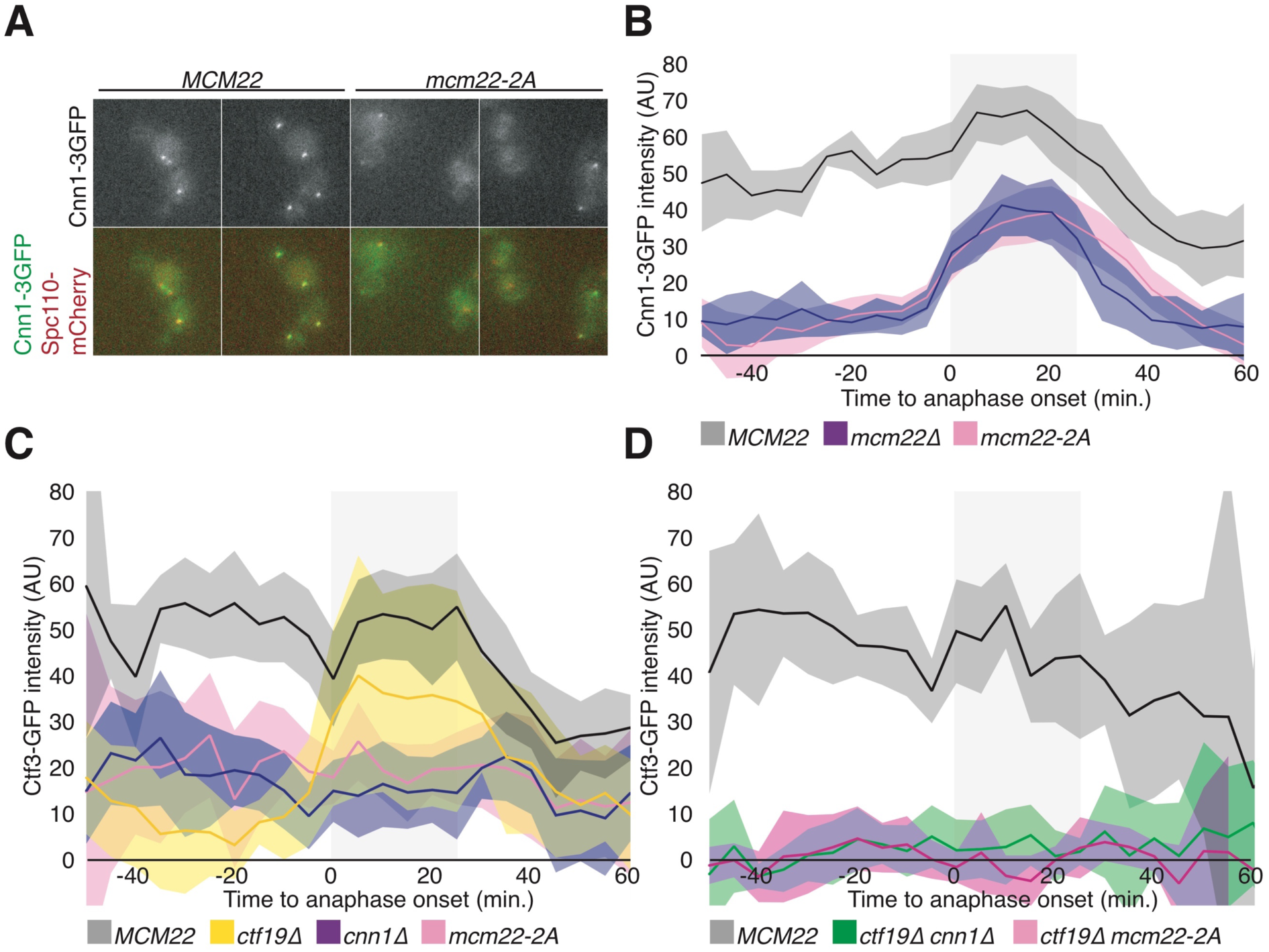
Live-cell imaging of Cnn1 and Ctf3. **(A)** Representative micrographs showing cells expressing Cnn1-3GFP, Spc110-mCherry, and either Mcm22-WT or Mcm22-2A. The same cells are shown at successive times during a single imaging session. **(B)** Kinetochore-associated Cnn1-3GFP intensity in cells from the indicated strains (*MCM22* – SMH798; *mcm22Δ* – SMH805; *mcm22-2A* – SMH799). Shaded box denotes approximate duration of anaphase. Lines denote mean intensity values (shaded areas – 95% confidence intervals, >12 cells per curve). **(C)** Kinetochore-associated Ctf3-GFP intensity shown as in (B) (*MCM22* – SMH728; *ctf19Δ* – SMH767; *cnn1Δ* – SMH768; *mcm22-2A* – SMH749). **(D)** Kinetochore-associated Ctf3-GFP intensity shown as in (B) (*MCM22* – SMH728; *ctf19Δ cnn1Δ* – SMH783; *ctf19Δ mcm22-2A* – SMH784).

We also examined Ctf3-GFP localization using the same live-cell imaging method (**Figure S4E**). If the Ctf3c recruits Cnn1-Wip1 to kinetochores, as is the case in a strictly hierarchical Ctf19c assembly model, then mutations that perturb Ctf3c-Cnn1 interaction should not affect Ctf3-GFP localization. In wild-type cells, the Ctf3-GFP signal at kinetochores peaks at anaphase and then dissipates, although less dramatically than the Cnn1-3GFP signal (**Figure 4C**). Cells expressing Ctf3-GFP and *mcm22-2A* show diminished but not ablated Ctf3-GFP localization throughout the cell cycle, a phenotype indistinguishable from *cnn1Δ* cells. Therefore, the Ctf3c and Cnn1-Wip1 display reciprocally dependent localization: each requires the other for efficient recruitment to the kinetochore.

Residual Ctf3-GFP localization in *mcm22-2A* and *cnn1Δ* cells indicates that the Ctf3c contacts the kinetochore through a second interface. The structure of the Ctf19c shows that the Ctf3c contacts Ctf19-Mcm21 and Iml3, with the major contribution to Ctf3c localization coming from Ctf19-Mcm21 (Hinshaw et al., 2019; Hinshaw and Harrison, 2019). Ctf3-GFP localization in *ctf19Δ* cells is nearly undetectable until anaphase, when the Ctf3 kinetochore signal is comparable to that seen in wild type cells (**Figure 4C**) (Pot et al., 2003). This signal complements the one seen in *mcm22-2A* and *cnn1Δ* cells, suggesting there are two collaborating pathways for Ctf3c recruitment: one through Ctf19-Mcm21 and the other through Cnn1-Wip1. Indeed, in cells lacking both pathways (*ctf19Δ cnn1Δ* or *ctf19Δ mcm22-2A*), the Ctf3-GFP signal is completely absent from kinetochores (**Figure 4D**). Therefore, two pathways contribute to Ctf3c kinetochore localization, and the Cnn1-dependent pathway dominates during anaphase.

## DISCUSSION

Kinetochore assembly proceeds in stages according to the cell cycle. In vertebrates, CDK1 and PLK1 determine the timing of CENP-A nucleosome deposition (McKinley and Cheeseman, 2014; Silva et al., 2012). Ipl1/Aurora B kinase determines the timing and extent of Ndc80 complex recruitment (Akiyoshi et al., 2013; Kim and Yu, 2015), which enables the outer kinetochore-microtubule connection. Ctf19c/CCAN factors link these regulated connections. We have determined the structure of the outermost Ctf19c components: the Ctf3c and Cnn1-Wip1. Using mutations in Mcm22, we have shown that Ctf19c assembly is also subject to cell cycle-dependent regulation. This regulation circumvents an observed hierarchical assembly pathway (Pekgoz Altunkaya et al., 2016), and its effects are most pronounced in anaphase.

The structure of the complete Ctf3c-Cnn1-Wip1 assembly enables unambiguous placement of Cnn1-Wip1 relative to other Ctf19c components (**Figure 5A**). Poorly resolved regions of a previous Ctf19c cryo-EM map (Hinshaw and Harrison, 2019) accommodate the model presented here, with a 31.4 Å displacement of the Ctf3-N module at its tip (measured from Mcm22-D168, **Figure S5A**). The Mcm16/22 linker region (**Figure 2C**) is the joint for the displacement. Cnn1-N, which is heavily phosphorylated *in vivo* (Bock et al., 2012; Malvezzi et al., 2013), projects towards the N-terminal helical domain of Ame1-Okp1 (analogous to MIND Head II (Dimitrova et al., 2016; Hinshaw and Harrison, 2019)), implying a possible regulated contact. That the disordered N-terminal regions of all three proteins cluster (**Figure S5B**) suggests coordinated targeting by mitotic kinases. We previously observed that the Ctf3-N module must swing outward to accommodate Cse4/CENP-A nucleosome contact (Hinshaw et al., 2019). The joint region in the current model provides a mechanism for this flexing, and the proximity of Ame1-N, Okp1-N, and Cnn1-N suggests that the flexing may be regulated.

**Figure 5.**
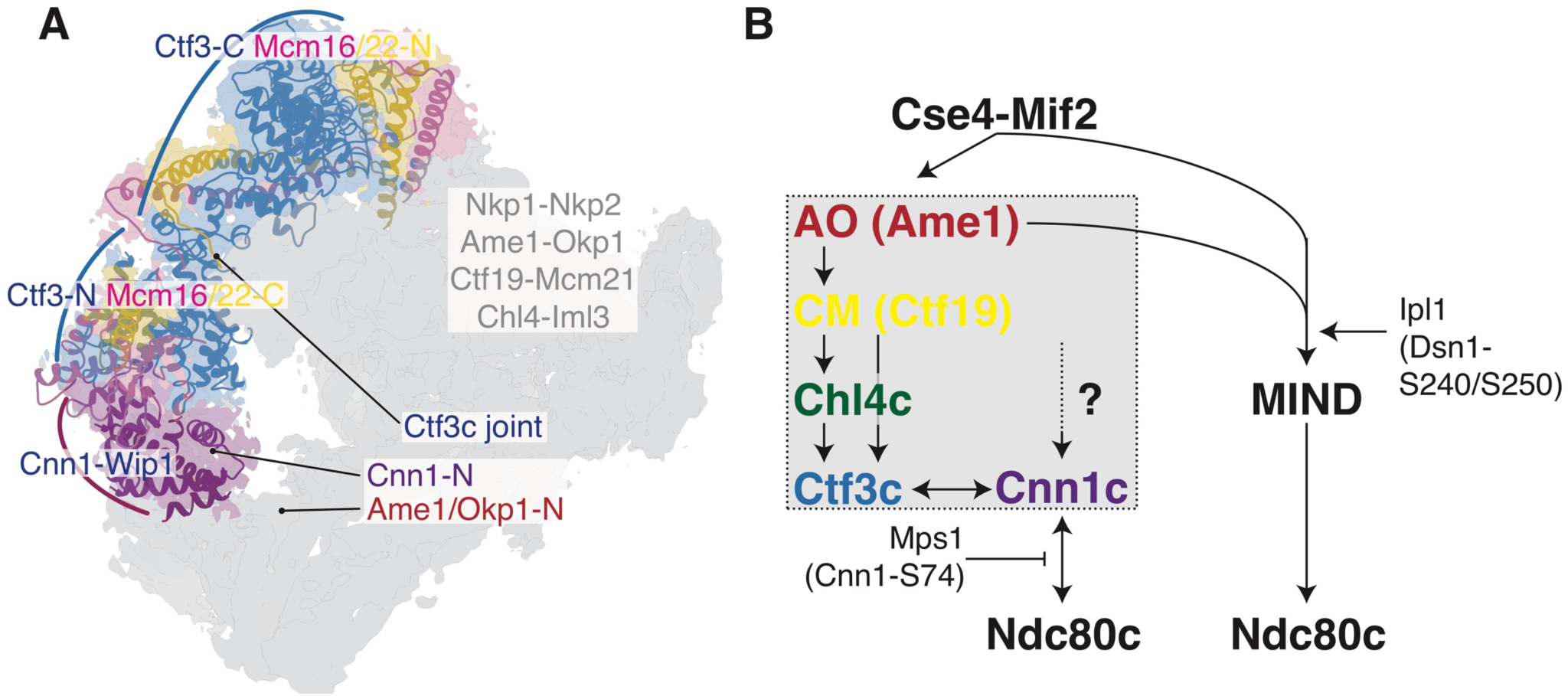
Ctf19c assembly model. **(A)** Updated three-dimensional model of the Ctf19c showing newly determined positions of the Ctf3-N, Mcm16/22-C, Cnn1, and Wip1. Partially transparent cryo-EM density for a single Ctf19c protomer is shown. The Ctf3c model was allowed to flex about its joint, bringing the Ctf3-N module (including Cnn1-Wip1) inward towards the Ame1/Okp1-N peptides (see **Figure S5A**). The Ctf3c joint and anchor points for flexible N-terminal extensions of Cnn1, Ame1, and Okp1 are indicated. **(B)** An updated model for hierarchical Ctf19c assembly showing that the Ctf3c and Cnn1-Wip1 depend on each other for localization, and Cnn1-Wip1 depends on Ndc80c interaction. The MIND-dependent pathway for centromere-microtubule contact is also shown. Ipl1 phosphorylates Dsn1, and Mps1 phosphorylates Cnn1 (AO – Ame1-Okp1, CM – Ctf19-Mcm21, Chl4c – Chl4-Iml3, Ctf3c – Ctf3-Mcm16-Mcm22, Cnn1c – Cnn1-Wip1, MIND – Mtw1-Nntf1-Nsl1-Dsn1, Ndc80c – Ndc80-Nuf2-Spc24-Spc25).

Although cryo-EM reconstructions show that Ctf19c proteins interdigitate, live-cell imaging experiments show that the kinetochore loading of some of these proteins varies during the cell cycle (Bock et al., 2012; Dhatchinamoorthy et al., 2017). This behavior is particularly pronounced for Cnn1; its recruitment depends on Ctf19c factors and is inhibited by Mps1 activity (Bock et al., 2012; Schleiffer et al., 2012; Thapa et al., 2015). We recapitulated these observations using live-cell imaging and found that two pathways contribute to Cnn1 localization. Precise inactivation of the Ctf19c-dependent pathway shows that the two pathways can operate independently. That the second pathway is only active during anaphase, when Mps1 is inactive (Palframan et al., 2006), strongly suggests it requires Cnn1-Ndc80c contact (Thapa et al., 2015). These observations imply complex feedback regulation: Mps1 activity inhibits Ctf19c-independent Cnn1 recruitment, and our findings show that this inhibition destabilizes Ctf3c kinetochore loading, further impairing Cnn1 recruitment. In vertebrate cells, Cnn1 binds the four-protein MIND complex upon phosphorylation by CDK1 (Huis In ‘t Veld et al., 2016; Rago et al., 2015). Although the CDK1 site is not obviously conserved in yeast, we cannot rule out that this or a related mechanism also contributes to Cnn1 localization. Restriction of Ctf19c-independent Cnn1 localization to anaphase suggests any such contribution must be Cdk1-independent. It remains to be seen how this regulatory logic has been modified in organisms that rely solely on the Cnn1/CENP-T pathway for kinetochore-microtubule contact (Cortes-Silva et al., 2020).

Accumulated evidence suggests MIND-Ndc80c and Cnn1-Ndc80c are parallel connections between centromeric DNA and spindle microtubules (**Figure 5B**) (Hara and Fukagawa, 2020). In vertebrate cells, mitotic kinases render the CENP-T connection dominant during anaphase (Hara et al., 2018). CENP-A misincorporation is not sufficient for CENP-H recruitment to an ectopic chromosomal locus (Gascoigne et al., 2011), implying centromere-specific regulation required for CCAN assembly. CENP-H and CENP-T are mutually required for robust localization (Hori et al., 2008). Similarly, our observation of Cnn1-dependent Ctf3 localization demonstrates that regulated outer kinetochore assembly indirectly influences the stability of the Ctf19c and that in yeast, as in vertebrate cells, Cnn1-mediated kinetochore stabilization is most pronounced during anaphase.

## ACKNOWLEDGMENTS

We thank the staff at the Harvard Cryo-Electron Microscopy Center for Structural Biology for help with high-resolution cryo-EM data collection, particularly Richard Walsh and Sarah Sterling. We thank Shaun Rawson for real-time movie processing. We thank SBGrid for computational support, particularly Mick Timony and Justin O’Connor. We thank Jennifer Waters and the staff at the Nikon Imaging Center at Harvard Medical School for light microscopy support. We thank Adèle Marston and Huilin Zhou for comments on the manuscript. S.M.H. is an HHMI fellow of the Helen Hay Whitney Foundation. S.C.H. is an Investigator of the Howard Hughes Medical Institute.

## AUTHOR CONTRIBUTIONS

S.M.H. conceived the study, performed the experiments, and wrote and edited the manuscript. S.C.H. supervised the work and edited the manuscript.

## FIGURES

**Figure S1.**
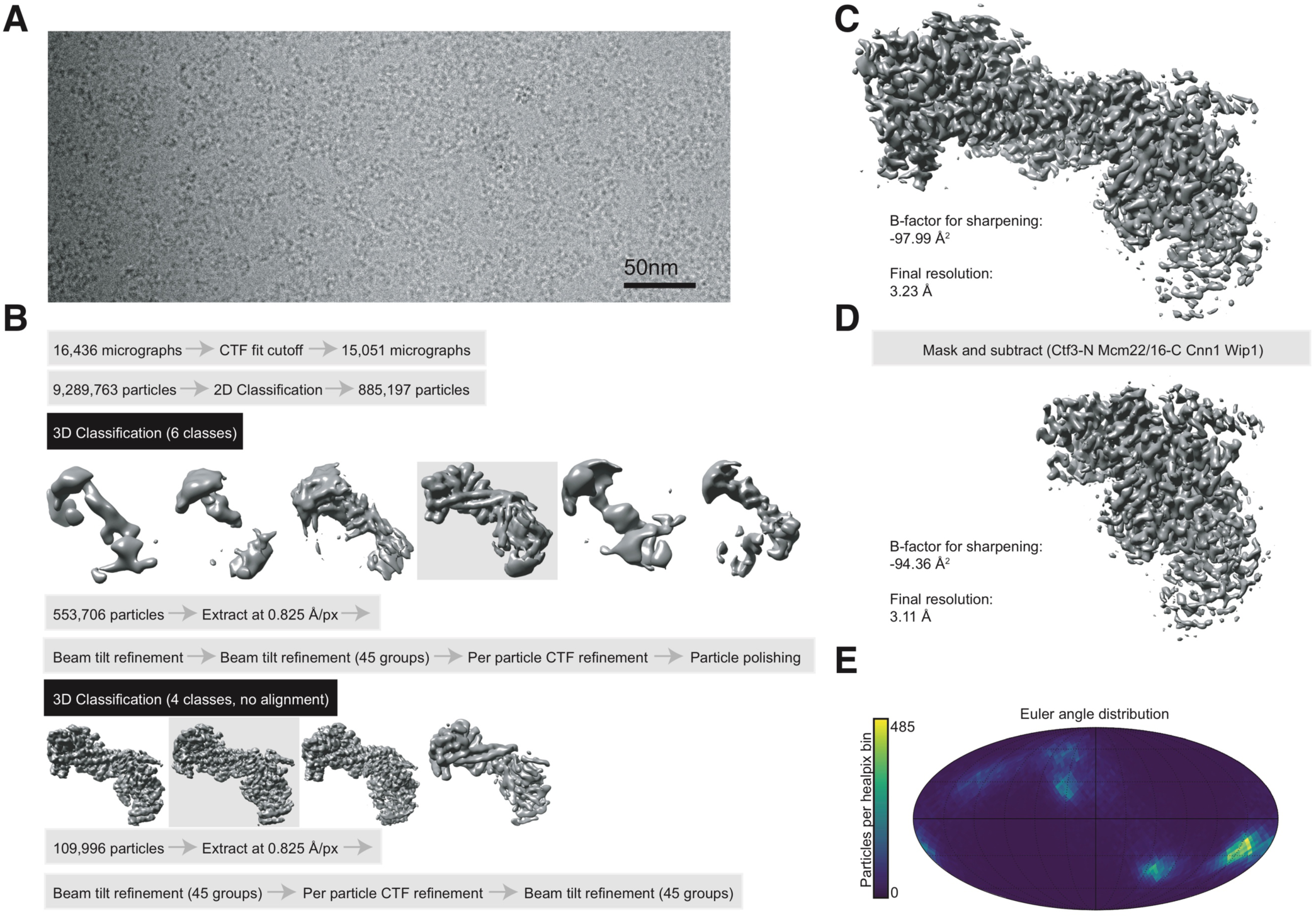
Data collection and cryo-EM map determination. **(A)** Representative cryo-electron micrograph. **(B)** Data processing steps for determination of the Ctf3c-Cnn1-Wip1 structure. **(C)** Final refined and B-factor sharpened map showing the Ctf3c-Cnn1-Wip1 complex. **(D)** Sharpened cryo-EM map for the Ctf3-N module including Cnn1-Wip1. **(E)** Euler angle distribution plot for the reconstruction shown in (C).

**Figure S2.**
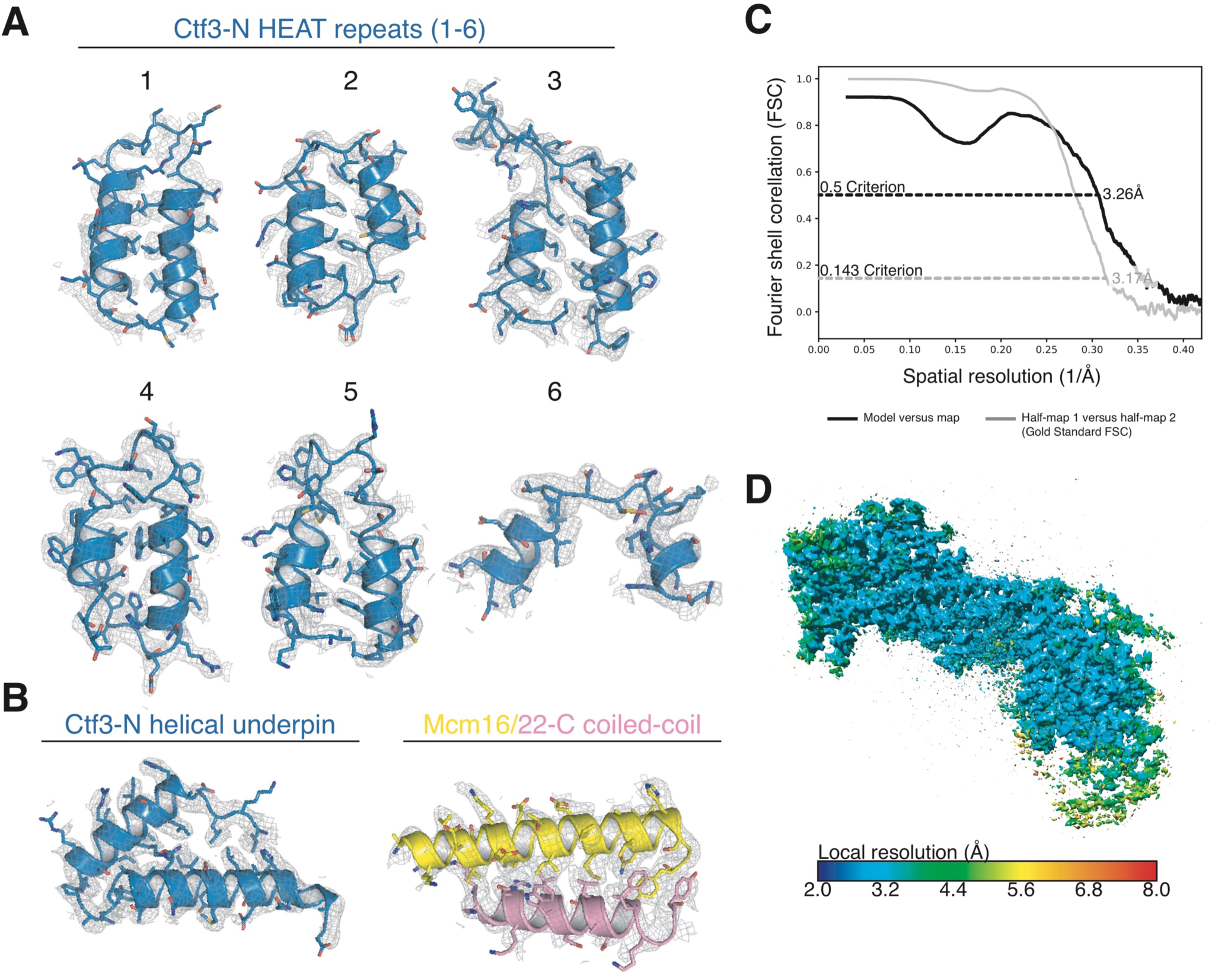
Molecular model quality. **(A)** Cryo-EM map and refined model showing the Ctf3-N HEAT repeats (pictured Ctf3 segments: 1 – 1-38; 2 – 40-74; 3 – 77-119; 4 – 121-159; 5 – 163-199; 6 – 201-223). **(B)** Cryo-EM map and refined density showing the Ctf3-N helical underpinning motif (Ctf3 224-268; left) and the Mcm16/22 C-terminal coiled-coil (Mcm16 148-174; Mcm22 141-159; right). **(C)** Fourier shell correlation curves describing the Ctf3c-Cnn1-Wip1 density map (half-map to half-map) and map-to-model correlations. **(D)** The refined cryo-EM map colored according to local resolution.

**Figure S3.**
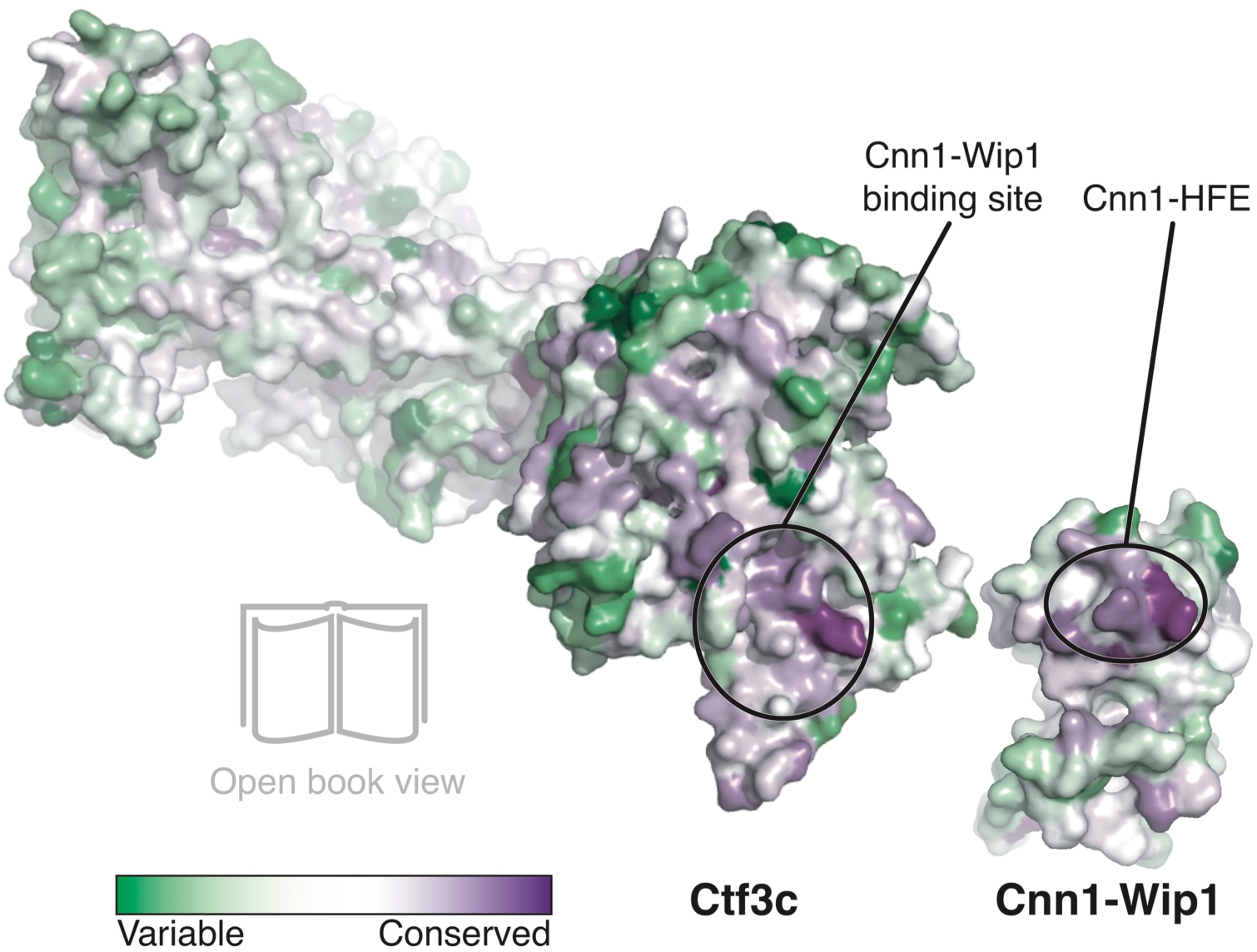
Conservation of the Ctf3c-Cnn1-Wip1 interface. The Ctf3c and Cnn1-Wip1 shown as space-filling models colored according to amino acid conservation across eukaryotes (Ashkenazy et al., 2016; van Hooff et al., 2017). Ctf3c and Cnn1-Wip1 models are split open to show their interaction surfaces (circled).

**Figure S4.**
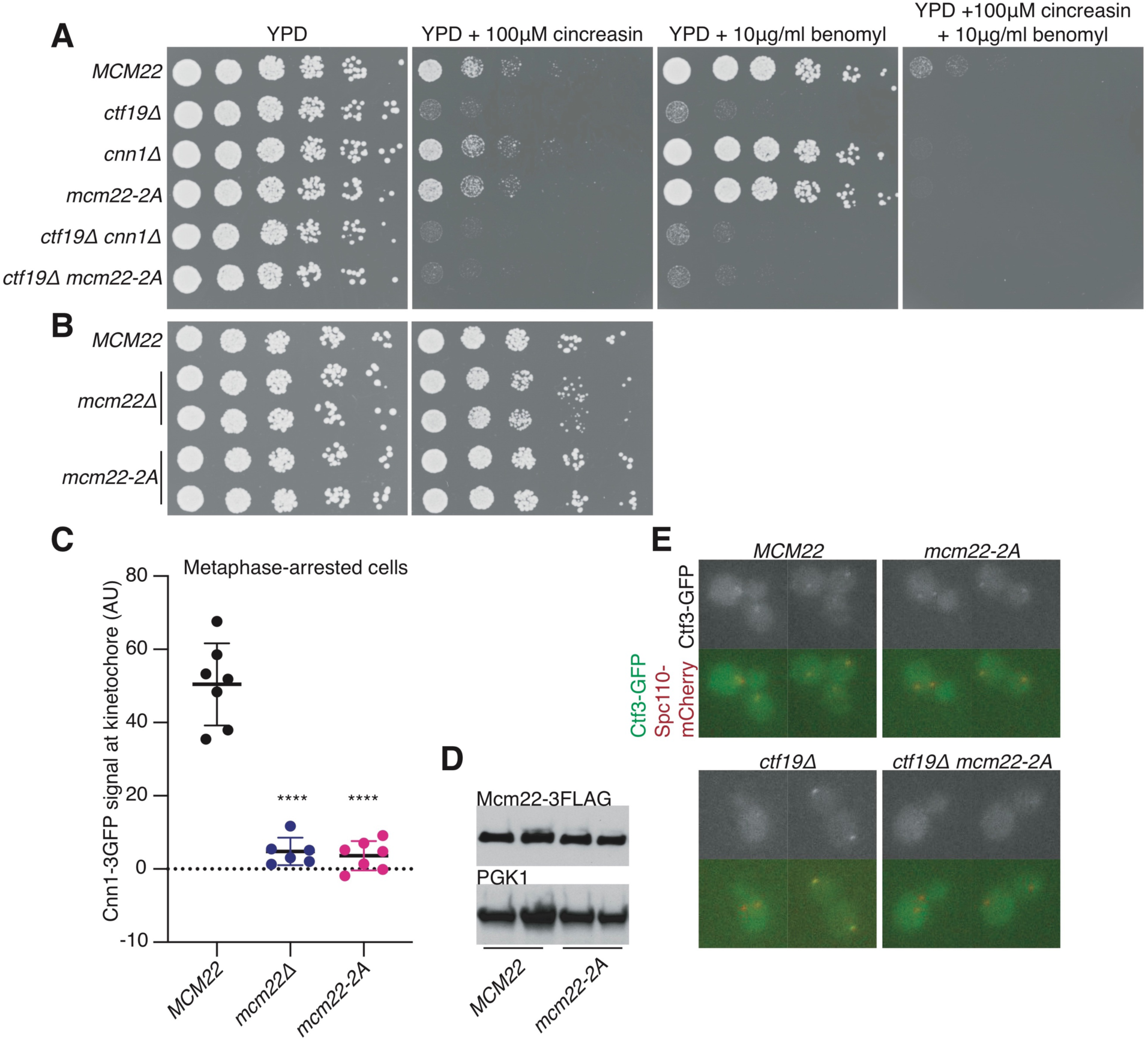
Phenotypes associated with mcm22-2A and related mutant strains. **(A)** Serial dilutions of the indicated strains grown on rich medium with the indicated additives (*MCM22* – SMH728; *ctf19Δ* – SMH767; *cnn1Δ* – SMH768; *mcm22-2A* – SMH749; *ctf19Δ cnn1Δ* – SMH783; *ctf19Δ mcm22-2A* – SMH784). **(B)** Serial dilutions of the indicated strains as in A (*MCM22* – SMH798; *mcm22Δ* – SMH805; *mcm22Δ* – SMH809; *mcm22-2A* – SMH799; *mcm22-2A* – SMH800). **(C)** Cnn1-3GFP intensity at kinetochores in metaphase-arrested cells (nocodazole and benomyl) of the indicated genotypes. Strains are from Figure 4B. At least six cells were quantified per group. Z-projected maximum intensity was averaged across six successive timepoints for each cell (**** – p < 0.0001, t-test, two tails, unequal variance). **(D)** Western blot showing Mcm22 protein levels for asynchronous cultures of the indicated genotypes (*MCM22* – SMH728 and SMH746; *mcm22-2A* – SMH749). **(E)** Representative micrographs showing Ctf3-GFP for the indicated genotypes (strains from panel A). Images presented as in Figure 4A.

**Figure S5.**
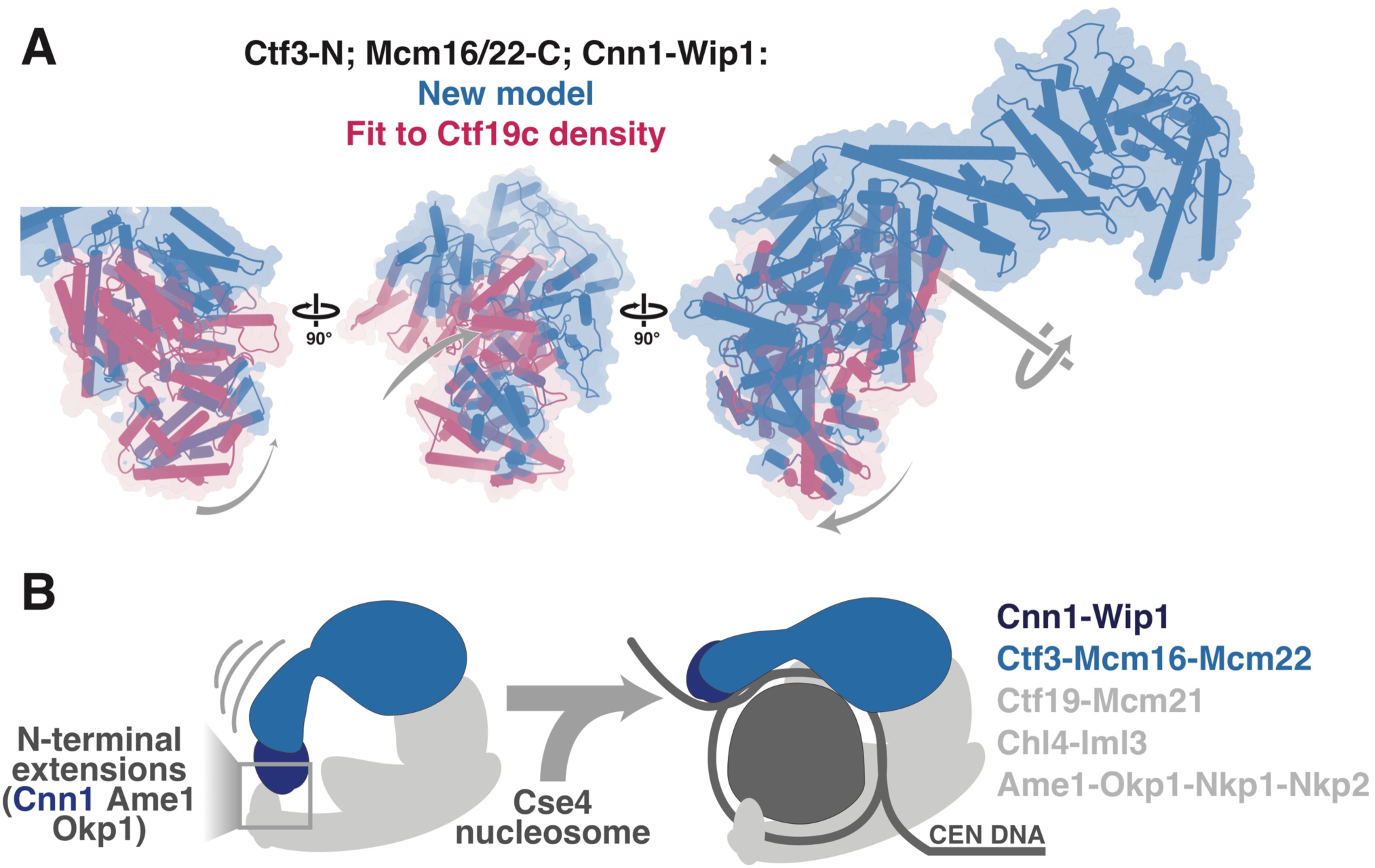
Displacement of the Ctf3-N module. **(A)** Model for the Cnf3-N module (Ctf3-N, Mcm16/22-C, Cnn1, Wip1) fit to the current cryo-EM map (blue) or the map of the full Ctf19c (pink) with the Ctf3-C module (Ctf3-C, Mcm16/22-N) held in place for reference. Three rotated views are shown. The gray arrows show movement of the Ctf3-N module from the Ctf19c density to the current map (Ctf3c-Cnn1-Wip1). The gray line (right) shows the axis of rotation about the Ctf3c joint. **(B)** Cartoon model showing predicted Ctf3-N module displacement required for accommodation of the Cse4/CENP-A nucleosome. The gray box shows where Cnn1-N, Ame1-N, and Okp1-N project from the well-ordered parts of the structure. Figure adapted from (Hinshaw et al., 2019).

## METHODS

### Resources Table

| Reagent type (species) or resource | Designation | Source or reference | Identifiers | Additional information |
| --- | --- | --- | --- | --- |
| Gene (*S. cerevisiae*) | See Table 4 |  |  |  |
| Strain, strain background (*S. cerevisiae*) | S288c |  |  |  |
| Strain, strain background (*S. cerevisiae*) | W303 |  |  |  |
| Genetic reagent (*S. cerevisiae*) | See Table 4 |  |  |  |
| Antibody | anti-FLAG-HRP (from ascites) | Abcam | 49763 | (1:1000) |
| Antibody | anti-PGK1 (mouse monoclonal) | Invitrogen | 459250 | (1:5000) |
| Antibody | goat anti-mouse-IgG-HRP (rabbit polyclonal) | Abcam | 97046 | (1:10000) |
| Recombinant DNA reagent | See Table 5 |  |  |  |
| Cell line (*E. coli*) | Rosetta 2(DE3)pLysS; *E. coli* | EMD Millipore | 71403 |  |
| Cell line (*T. ni*) | High Five cells; *Trichoplusia ni* | ThermoFisher | B85502 | Harrison lab stock |
| Software, algorithm | SerialEM (v3.7) | (Mastronarde, 2005) |  |  |
| Software, algorithm | MotionCor2 (v1.1.0) | (Zheng et al., 2017) |  |  |
| Software, algorithm | CTFFIND4 (v4.1.8) | (Rohou and Grigorieff, 2015) |  |  |
| Software, algorithm | Relion (v3.0.1) | (Zivanov et al., 2018) |  |  |
| Software, algorithm | ResMap (v1.1.4) | (Kucukelbir et al., 2014) |  |  |
| Software, algorithm | PyMol (v2.3.2) | Schrödinger, LLC |  |  |
| Software, algorithm | Chimera (v1.14.0) | (Pettersen et al., 2004) |  |  |
| Software, algorithm | Coot (v0.8.8) | (Emsley et al., 2010) |  |  |
| Software, algorithm | Phenix (v1.17.1) | (Afonine et al., 2018) |  |  |
| Software, algorithm | MAFFT | (Katoh et al., 2019) |  |  |
| Software, algorithm | JalView | (Waterhouse et al., 2009) |  |  |
| Software, algorithm | SWISS-MODEL | (Waterhouse et al., 2018) |  |  |
| Software, algorithm | Fiji | (Schindelin et al., 2012) |  |  |
| Software, algorithm | python 2.7.2 | www.python.org |  |  |
| Software, algorithm | ConSurf server | (Ashkenazy et al., 2016) |  |  |
| Software, algorithm | Pirsm v8.4.1 | GraphPad |  |  |
| Other (holey carbon grids) | C-flat | Electron Microscopy Sciences | CF-2/1-4C |  |

### Protein purification and cryo-EM sample preparation

Recombinant kinetochore proteins were purified as described previously and stored as aliquots at -80 degrees Celsius (Hinshaw and Harrison, 2019). To reconstitute the sample used for cryo-EM, equimolar amounts of purified Ctf3c and Cnn1-Wip1 were mixed and incubated on ice for one hour before injection onto a size exclusion column (Superdex 200 Increase 5/150 GL) developed in gel filtration buffer (20 mM Tris-HCl, pH 8.5; 150 mM NaCl, 1 mM EDTA; 1 mM TCEP; 0.02% Na-azide (*w:v*)). The sample was eluted by isocratic flow at 0.1 mL/min, and fractions were collected at 30 second intervals. For the sample reported here, excess recombinant NH_2_-biotin-Ulp2-CCR peptide (Suhandynata et al., 2019) was included but not visualized in the final density.

After verification of complex formation by SDS-PAGE, the purified sample was vitrified for cryo-EM. 3.5 µL of gel filtration column eluate (∼0.5 mg/ml) was applied to a holey carbon support (Quantifoil 2/1-4Cu-50) at ∼90% relative humidity. The grid was immediately blotted on both sides for four seconds and plunged into liquid ethane (Gatan CryoPlunge 3).

### Cryo-EM data collection

Data collection was done as described previously (Hinshaw et al., 2019) using SerialEM (Mastronarde, 2005). A Titan Krios G3i (Gatan) operating at 300 kV was used to illuminate the sample with a 1 µm beam in nanoprobe mode with a pre-camera energy filter (Gatan) slit width of 100 eV. Movies were collected on a K3 detector (Gatan) in counting mode with a pixel size of 0.825 µm. 50 frames were collected per 3 second movie, with a total dose of 60 electrons/Å^2^ divided equally among the frames. The defocus range was set between -1.2 and -3.0 µm. Using image shift and real-time coma-correction by beam tilt as implemented in SerialEM, nine holes were visited at each stage position, and five movies were taken per hole, giving a total of 45 movies per stage movement.

### Cryo-EM data processing and map determination

Initial movie processing steps were carried out using an in-house pipeline (Shaun Rawson) built upon RELION-3.0 software (Zivanov et al., 2018). Movies, stored as LZW-compressed tiffs, were aligned using MotionCorr2 software (5-by-5 patches) (Zheng et al., 2017), and CTFFIND4 was used for CTF parameter estimation (Rohou and Grigorieff, 2015). All subsequent processing steps were carried out in Relion 3.0 after data collection. Micrographs were discarded if the maximum resolution estimation (CTFFIND4) was worse than 4 Å. Data processing steps are given in detail in Figure S1. Care was taken after 2D-classification to select only projections showing features corresponding to the Ctf3-N module, and this proved essential for generating a high-quality map of the complete complex. Map quality and mask appropriateness were assessed visually in Chimera. The ResMap program was used to estimate local resolution throughout the map (Kucukelbir et al., 2014).

### Molecular modeling

Deposited coordinates describing Ctf3-C module (PDB: 6OUA) (Hinshaw et al., 2019) was used as a starting model for the corresponding part of the density. Minor modifications were required, including updates to disordered segments between HEAT repeats and parts of the Ctf3 helical knob. The C-terminal-most alpha helical segment of Ctf3 is also well-ordered in the current model, whereas it was poorly represented in the previous map. We used SWISS-MODEL (Waterhouse et al., 2018) to generate a homology model of *S. cerevisiae* Ctf3-N from a crystal structure of this domain from *Chaetomium thermophilum* (PDB: 5Z07) (Hu et al., 2019). Mcm16/22-C and their connections to Mcm16/22-N were built *de novo*. A homology model for the Cnn1-Wip1 histone fold domain was constructed similarly using a crystal structure of the chicken homologs (PDB: 3B0C) (Nishino et al., 2012). The homology models were docked into the cryo-EM density using Chimera v1.14.0 and modified using COOT v0.8.8. The map showing the Cff3-N module was used for adjusting the corresponding atomic coordinates in later stages of model building. Phenix v1.17.1 Real Space Refine (Afonine et al., 2018) was used as a final step to optimize model geometry. All automated refinements and model statistics calculations were done using the map of the full complex. The Phenix component Mtriage was used to compute the model parameters, its fit to the density, and to generate a final resolution measurement from half-maps.

For analysis of amino acid conservation, multiple sequence alignments were created with MAFFT and displayed with JalView (Katoh et al., 2019; Waterhouse et al., 2009). Surface rendering showing amino acid conservation was generated by the ConSurf server (Ashkenazy et al., 2016). Inputs for multiple sequence alignments were taken from van Hooff and modified if necessary (van Hooff et al., 2017).

### Yeast strain generation and growth

Yeast strains were generated by standard methods and grown in rich medium with additives or dropout medium (synthetic complete (SC), Sunrise Science Products) as required. Cincreasin (6-bromo-1,3-benzoxazol-2(3h)-one, Fisher Scientific) was diluted to 100 mM in DMSO immediately before use as described previously (Dorer et al., 2005). Benomyl (10 mg/mL stock solution in DMSO) was mixed with near-boiling medium to prevent crystallization. For metaphase arrests, cells were grown as for other imaging experiments (below) and arrested by addition of 10 µg/ml benomyl and 15 µg/ml nocodazole. After one hour, benomyl was re-added at the same concentration. Cells were imaged as described below two hours after the arrest was initiated.

The Cnn1-3GFP Spc110-mCherry strain was a generous gift from Peter De Wulf (Bock et al., 2012). Derivatives were generated by lithium acetate-mediated integration of PCR products (Longtine et al., 1998). Ctf3-GFP Spc110-mCherry strains were generated similarly, except *ctf19Δ mcm22-2A* and *ctf19d cnn1Δ* strains were created by mating of the individual mutants in the imaging background and subsequent sporulation. Western blotting was carried out as described (Hinshaw et al., 2019) with the following antibodies: anti-FLAG-HRP (Abcam 49763 from ascites), anti-PGK1 (Invitrogen 459250), and rabbit anti-mouse IgG-HRP (Abcam 97046).

### Fluorescence imaging and quantification

Saturated overnight cultures of the indicated strains were grown in SCA (SC supplemented with 20 µg/ml additional adenine). Cultures were diluted 1:50 (*v:v*) into new SCA and grown at least four hours longer before mounting on concanavalin A-coated cover slips for imaging. Cover slips were mounted in a humidified chamber heated to ∼29 degrees Celsius using a Tokai Hit stage-top incubator.

Images were taken with NIS-Elements acquisition software used to run a Nikon Ti2 motorized inverted microscope with the Perfect Focus System, a Lumencor SpectraX fluorescence light source, a Nikon Plan Apo 60x NA objective, and a Hamamatsu Flash4.0 V2+ sCMOS camera. Optical paths were: GFP – SpectraX Cyan (illumination), Lumencor 470/24 (excitation), and Semrock FF03 525/50 (emission); mCherry – SpectraX GreenYellow (illumination), Lumencor 525/25 (excitation), and Semrock FF02 641/75 (emission). Nine z-heights (0.4 µm spacing) were taken per stage position. Timepoints were taken at five-minute intervals for at least two hours for each imaging session.

Maximum z-projections for each timepoint were analyzed for five stage positions for each indicated strain. To get intensity values, a 6 pixel circle was drawn around a single kinetochore cluster, and background signal in an adjacent nuclear region was subtracted for each timepoint using Fiji (Schindelin et al., 2012). Irreversible Spc110-mCherry separation was taken as an indication of anaphase initiation. *ctf19Δ* cells show pronounced and prolonged Spc110 separation before anaphase initiation due to inefficient sister centromere cohesion, and care was taken to differentiate this from *bona fide* anaphase initiation. At least 12 cells were included for each measurement displayed. All traces shown on the same plot originated from the same experiment. Plots were generated with GraphPad Prism v8.4.1 software (GraphPad). Experiments were repeated at least twice for the indicated genotypes with equivalent results. Imaging conditions were kept constant for all experiments, although the GFP illumination time differed for Ctf3-GFP and Cnn1-3GFP strains.

